# Gene editing of *SAMHD1* in macrophage-like cells reveals complex relationships between SAMHD1 phospho-regulation, HIV-1 restriction and cellular dNTP levels

**DOI:** 10.1101/2023.08.24.554731

**Authors:** Moritz Schüssler, Kerstin Schott, Nina Verena Fuchs, Adrian Oo, Morssal Zahadi, Paula Rauch, Baek Kim, Renate König

## Abstract

Sterile α motif (SAM) and HD domain-containing protein 1 (SAMHD1) is a dNTP triphosphate triphosphohydrolase (dNTPase) and a potent restriction factor for immunodeficiency virus 1 (HIV-1), active in myeloid and resting CD4^+^ T cells. The anti-viral activity of SAMHD1 is regulated by dephosphorylation of the residue T592. However, the impact of T592 phosphorylation on dNTPase activity is still under debate. Whether additional cellular functions of SAMHD1 impact anti-viral restriction is not completely understood.

We report BLaER1 cells as a novel human macrophage HIV-1 infection model combined with CRISPR/Cas9 knock-in (KI) introducing specific mutations into the *SAMHD1* locus to study mutations in a physiological context. Transdifferentiated BLaER1 cells harbor active dephosphorylated SAMHD1 that blocks HIV-1 reporter virus infection. As expected, homozygous T592E mutation, but not T592A, relieved a block to HIV-1 reverse transcription. Co-delivery of VLP-Vpx to SAMHD1 T592E KI mutant cells did not further enhance HIV-1 infection indicating the absence of an additional SAMHD1-mediated antiviral activity independent of T592 de-phosphorylation. T592E KI cells retained dNTP levels similar to WT cells indicating uncoupling of anti-viral and dNTPase activity of SAMHD1. The integrity of the catalytic site in SAMHD1 was critical for anti-viral activity, yet poor correlation of HIV-1 restriction and global cellular dNTP levels was observed in cells harboring catalytic core mutations. Together, we emphasize the complexity of the relationship between HIV-1 restriction, SAMHD1 enzymatic function and T592 phospho-regulation and provide novel tools for investigation in an endogenous and physiological context.

**Importance:** We introduce BLaER1 cells as an alternative myeloid cell model in combination with CRISPR/Cas9-mediated gene editing to study the influence of SAMHD1 T592 Mophosphorylation on anti-viral restriction and the control of cellular dNTP levels in an endogenous, physiological relevant context. Proper understanding of the mechanism of the anti-viral function of SAMHD1 will provide attractive strategies aiming at selectively manipulating SAMHD1 without affecting other cellular functions.

Even more, our toolkit may inspire further genetic analysis and investigation of restriction factors inhibiting retroviruses, their cellular function and regulation, leading to a deeper understanding of intrinsic anti-viral immunity.

## Introduction

Sterile α motif (SAM) and HD domain-containing protein 1 (SAMHD1) is a potent anti-viral restriction factor with broad anti-viral activity against a number of viruses, including lenti- and non-lenti retroviruses (for review see (Majer et al. 2019)). In particular, HIV-1 replication is restricted in myeloid cells and resting CD4^+^ T cells (Laguette et al. 2011; Hrecka et al. 2011; Berger et al. 2011; Baldauf et al. 2012; Descours et al. 2012). SAMHD1 depletion leads to an increase in intermediates of reverse transcription (RT), especially late cDNA products, indicating that SAMHD1 inhibits the RT process (Hrecka et al. 2011; Baldauf et al. 2017; Schott et al. 2018).

SAMHD1 is a cellular dNTP triphosphate triphosphohydrolase (dNTPase). It is active as a tetramer, regulated by binding of GTP/dGTP and dNTPs to primary and secondary allosteric sites, respectively (Hansen et al. 2014). Therefore, the obvious assumption might be that SAMHD1 inhibits HIV-1 replication through depletion of dNTPs, the substrate for HIV-1 RT (Goldstone et al. 2011; Lahouassa et al. 2012). Providing exogenous desoxyribonucleotides (dNs) rescues HIV-1 replication in cells expressing SAMHD1 (Baldauf et al. 2012; Lahouassa et al. 2012). In addition, SAMHD1 mutants shown to lack dNTPase activity, both *in vitro* or in cells, lose their restrictive potential, when overexpressed in phorbol 12-myristate-13-acetate (PMA)-activated macrophage-like U937 cells (Laguette et al. 2011; Lahouassa et al. 2012; Arnold et al. 2015; White et al. 2013a). However, SAMHD1 dNTPase activity might not be sufficient for HIV-1 restriction (Majer et al. 2019; Welbourn und Strebel 2016). It is hypothesized that additional SAMHD1 mediated functions like modulation of immune signaling, resolution of stalled replication forks and R-loops, RNA binding, or its role in DNA damage response, might contribute to the restrictive phenotype (Majer et al. 2019; Chen et al. 2018; Coquel et al. 2018; Park et al. 2021; Daddacha et al. 2017).

Only SAMHD1 dephosphorylated at residue T592 is active against HIV-1 (Cribier et al. 2013; White et al. 2013b; Welbourn et al. 2013). SAMHD1 is phosphorylated in cycling cells by cyclin dependent kinases CDK1 and CDK2 in complex with Cyclin A2 in S and G_2_/M phase (White et al. 2013b; Cribier et al. 2013). At mitotic exit, SAMHD1 is rapidly dephosphorylated at residue T592 due to the action of the PP2A-B55α phosphatase complex (Schott et al. 2018). While the effect of SAMHD1 T592 phosphorylation on HIV-1 restriction is consistently demonstrated, the consequence for its dNTPase activity is still under debate. Biochemical approaches to measure the effect of SAMHD1 T592 phosphorylation and phosphomimetic mutants on SAMHD1 tetramer formation and dNTPase activity have not been able to reveal a functional relationship (Arnold et al. 2015; White et al. 2013b; Welbourn et al. 2013; Bhattacharya et al. 2016; Yan et al. 2015). Still, cell cycle dependent SAMHD1 phosphorylation, loss of HIV-1 restriction and increased dNTP levels in S- and G_2_/M phase in synchronized HeLa cells show a clear timely correlation (Schott et al. 2018). In contrast, mutagenic analysis of T592 site in myeloid cells challenges a causative link. Phosphoablative T592A or T592V, but not phosphomimetic T592E or T592D mutants, were able to inhibit HIV-1 replication, when overexpressed in PMA activated U937 cells (Arnold et al. 2015; White et al. 2013b; Welbourn et al. 2013; Ryoo et al. 2014). Conversely, not only phosphoablative but also phosphomimetic SAMHD1 T592 mutants efficiently limited the cellular dNTP pool (Welbourn und Strebel 2016; White et al. 2013b; Welbourn et al. 2013). This obvious discrepancy might be due to biological reasons (for details refer to discussion and review (Majer et al. 2019)). However, also technical limitations might be the cause for this problem.

Genetic studies of SAMHD1 phospho-mutants in myeloid cells are currently limited to PMA-activated macrophage-like THP-1 or U937 cells. As treatment with PMA can activate non-physiological intracellular pathways (Chanput et al. 2010; Zeng et al. 2015), alternative myeloid models are needed, which ideally would be both genetically amendable and based on physiological myeloid differentiation pathways.

So far, anti-viral restriction has been tested with mutant constructs of SAMHD1 using lenti- or retroviral transduction. In this case, an exogenous promotor mediates overexpression of SAMHD1. The use of CRISPR/Cas9 allowed us to modify *SAMHD1* within the native genetic environment and to analyze the impact of selected mutations on anti-viral restriction in a physiological context, avoiding potential unwanted effects of mutant protein overexpression.

Here, we use CRISPR/Cas9-mediated knock-in (KI) in combination with transdifferentiated macrophage-like BLaER1 cells as a tool to study the impact of SAMHD1 T592 phosphorylation on HIV-1 restriction and dNTP pools in myeloid cells. Transdifferentiated macrophage-like BLaER1 cells expressed SAMHD1, which was dephosphorylated at residue T592. Concomitantly, transdifferentiated BLaER1 cells restricted HIV-1 replication in a SAMHD1 dependent manner. Introduction of SAMHD1 homozygous T592E mutations via CRISPR/Cas9 KI led to loss of HIV-1 restriction, while SAMHD1 T592A mutants maintained their anti-viral activity. Interestingly, HIV-1 infection was not further enhanced by SAMHD1 T592E mutant depletion suggesting the absence of an additional anti-viral activity of SAMHD1 that is independent of T592 de-phosphorylation. This highlights the T592 phospho-site as the critical residue for anti-viral activity. Remarkably, neither endogenous SAMHD1 T592E, nor T592A mutants, had an impact on cellular dNTP levels in transdifferentiated BLaER1 cells, indicating that the regulation of anti-viral and dNTPase activity of SAMHD1 can be uncoupled. However, mutagenic analysis of the catalytic residues H210, D218 and D311 highlights the importance of the integrity of the catalytic site for anti-viral restriction. Yet, we observed again a lack of correlation between cellular dNTP levels and HIV-1 restriction potential, indicating that the relationship between SAMHD1 function, HIV-1 restriction and T592 phospho-regulation is complex. Also we emphasize that regulation of dNTP levels is neither sufficient, nor necessary for SAMHD1-dependent HIV-1 restriction in macrophage-like BLaER1 cells.

## Results

### SAMHD1 is dephosphorylated at residue T592 in macrophage-like BLaER1 cells

Myeloid models to study HIV-1 restriction by mutagenesis are very limited. Transdifferentiated BLaER1 cells are a novel myeloid cell model, which has successfully been used to study innate immune signaling in macrophage-like cells (Gaidt et al. 2016; Rapino et al. 2013). The native, B-lineage derived BLaER1 cells undergo macrophage transdifferentiation by induction of the myeloid transcription factor C/EBPα. Transdifferentiated BLaER1 cells have been shown to closely resemble human macrophages with respect to mRNA expression, cell cycle arrest and immune functions (Rapino et al. 2013; Gaidt et al. 2018). In order to test whether these cells can serve as a model to study SAMHD1 mediated anti-viral restriction, we analyzed SAMHD1 expression in transdifferentiated BLaER1 cells. Flow cytometry analysis of transdifferentiated BLaER1 cells showed loss of B cell marker CD19 and acquisition of surface expression of the macrophage marker CD11b (Fig. 1A), as demonstrated earlier (Rapino et al. 2013). Transdifferentiation of BLaER1 cells using an adopted protocol, was highly reproducible and yielded 89.3 ± 8.8% (n = 33) of CD19^-^ CD11b^+^ cells in viable BLaER1 cells expressing GFP (Fig. 1B). In addition, transdifferentiated BLaER1 cells expressed monocyte-derived macrophage and dendritic cell markers CD14, CD163, CD206 and CD11c (Fig. 1C) highlighting the myeloid phenotype of these cells and validating previous results based on mRNA expression (Uhlen et al. 2019; Karlsson et al. 2021; Rapino et al. 2013). Interestingly, transdifferentiated BLaER1 cells displayed HIV-1 entry receptor CD4 expression and high surface level expression of both co-receptors CXCR4 and CCR5 indicating that they may be amenable to infection with CCR5- and CXCR4-tropic HIV-1 (Fig. 1C). Transdifferentiated BLaER1 cells expressed levels of SAMHD1 comparable to cycling THP-1 cells (Fig. 1D), whereas native BLaER1 cells showed no SAMHD1 expression. As T592 phosphorylation in SAMHD1 is the major regulator of anti-viral restriction (Cribier et al. 2013; White et al. 2013b; Schott et al. 2018), we analyzed the phosphorylation status in transdifferentiated BLaER1 cells. Relative SAMHD1 T592 phosphorylation was 31-fold lower in transdifferentiated BLaER1 compared to cycling THP-1 cells (0.032 ± 0.013 relative SAMHD1 pT592 normalized to cycling THP-1, n = 6) and in fact was barely detectable by immunoblotting even after long exposure times (Fig. 1D). Absence of SAMHD1 pT592 correlated well with the reported G_1_/G_0_ cell cycle arrest in transdifferentiated BLaER1 cells (Schott et al. 2018; Rapino et al. 2013), as well as with low cyclin A2 expression (Fig. 1D), which in complex with CDK1 and CDK2 is known to mediate T592 phosphorylation (Cribier et al. 2013). Thus, macrophage-like transdifferentiated BLaER1 cells expressed SAMHD1 dephosphorylated at residue T592, suggesting it to be anti-virally active.

**Figure 1:**
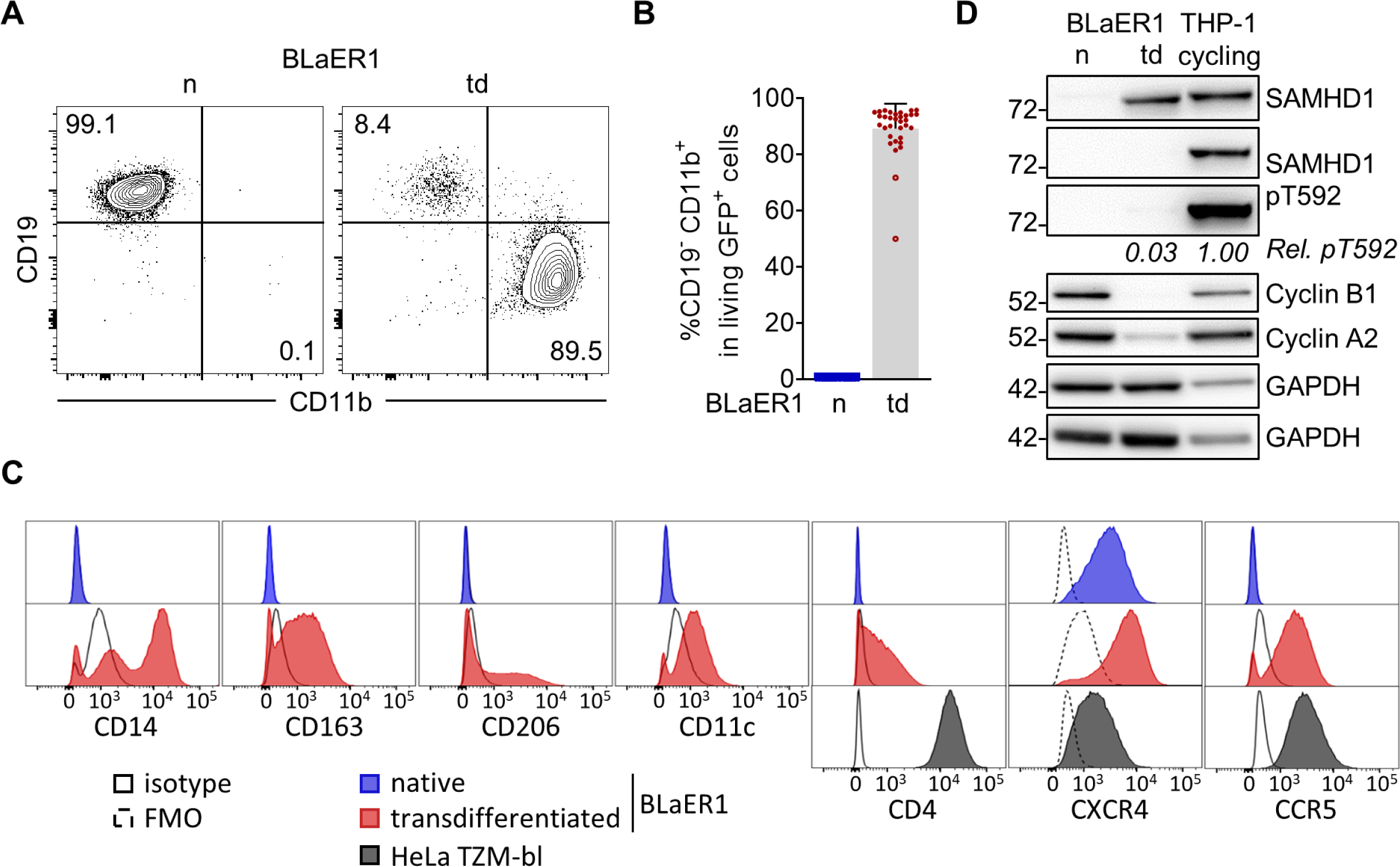
SAMHD1 is dephosphorylated at residue T592 in macrophage-like BLaER1 cells. **(A)** Representative flow cytometry analysis of CD19 and CD11b surface expression in native (n) and transdifferentiated (td) BLaER1 cells. Relative frequencies of CD19^+^ CD11b^-^ and CD19^-^ CD11b^+^ cell populations are indicated as % of viable GFP^+^ cells (n = 33). **(B)** Relative quantification of macrophage-like CD19^-^ CD11b^+^ cells in viable GFP^+^ native (n) or transdifferentiated (td) BLaER1 cells. Every dot represents an individual transdifferentiation approach. Experiments in which transdifferentiated BLaER1 cells show < 75% CD19^-^ CD11b^+^ cells in viable GFP^+^ cells were excluded from downstream analysis (open circles). Error bars represent standard deviation (n_n_ = 30, n_td_ = 33). **(C)** Surface expression of indicated monocyte-derived macrophage or dendritic cell associated markers CD14, CD163, CD206 or CD11c respectively, as well as HIV-1 (co-) receptors CD4, CXCR4 (CD184) and CCR5 (CD195), as analyzed by flow cytometry in viable GFP^+^ cells of native (blue) and transdifferentiated (red) BLaER1 cells. HeLa TZM-bl cells were used as positive controls for HIV-1 (co-) receptors. Solid or dashed black lines indicate respective isotype or fluorescence minus one (FMO) controls (n = 3). **(D)** Representative immunoblot analysis of SAMHD1, Cyclin B1 and Cyclin A2 expression in native (n) and transdifferentiated (td) BLaER1 cells, as well as cycling THP-1 cells. GAPDH serves as a loading control. Mean signal of SAMHD1 T592 phosphorylation (pT592) relative to total SAMHD1 expression in transdifferentiated BLaER1 cells was normalized to cycling THP-1 (n = 6).

### SAMHD1 restricts HIV-1 in macrophage-like BLaER1 cells

To define the restrictive capacity of SAMHD1 in the context of transdifferentiated BLaER1 cells, we infected the cells with a single-cycle HIV-1 luciferase reporter virus (HIV-1-luc), in presence or absence of virus like particles containing Vpx (VLP-Vpx). VLP-Vpx treatment led to efficient degradation of SAMHD1 (0.013 ± 0.007 relative SAMHD1 expression normalized to no VLP-Vpx, n = 3) (Fig. 2A) and increased HIV-1-luc infection. Linear regression revealed a significant (*p = 0.0125*, n = 3, unpaired t-test) increase over a wide range of MOIs (Fig. 2B). To validate this further, we generated SAMHD1 knock-out (KO) BLaER1 cells using CRISPR/Cas9 ribonucleoprotein (RNP). Three independent SAMHD1 KO BLaER1 single cell clones were analyzed in detail and showed bi-allelic InDels at the intended target site (Fig. 2C) leading to a frameshift, the introduction of premature stop codons and therefore, absence of SAMHD1 expression in transdifferentiated BLaER1 cells (Fig. 2D). While SAMHD1 KO did not affect BLaER1 transdifferentiation (Fig. 2E), it strongly increased HIV-1-luc infection at 24 hpi, as compared to wild type (WT) cells. Significance of differences in the slopes of linear regressions are suggesting SAMHD1 to be a major restriction factor in these cells over a wide range of MOIs (*p < 0.0001* for Clone #1, 2 and 3, n = 7, One-way ANOVA) (Fig. 2F). In order to rule out a potential confounding effect of a minor CD11b^-^ native-like population, we developed a flow cytometry workflow combining the use of a single-cycle HIV-1 mCherry (HIV-1-mCherry) reporter virus together with staining for viable CD11b^+^ cells. Thereby, we could specifically analyze infection in transdifferentiated CD11b^+^ macrophage-like BLaER1 cells. HIV-1-mCherry infection, as measured by %mCherry^+^ cells in CD11b^+^ viable GFP^+^ transdifferentiated BLaER1 cells, was strongly increased upon SAMHD1 KO at 24 hpi (Clone #1 *p = 0.3633*, #2 *p = 0.0360*, #3 *p = 0.0013*, n = 5, Kruskal–Wallis test) (Fig. 2G and H). This indicates that SAMHD1 is a major anti-lentiviral restriction factor in macrophage-like transdifferentiated BLaER1 cells. We therefore conclude that transdifferentiated BLaER1 cells are an excellent model to study SAMHD1 mediated HIV-1 restriction.

**Figure 2:**
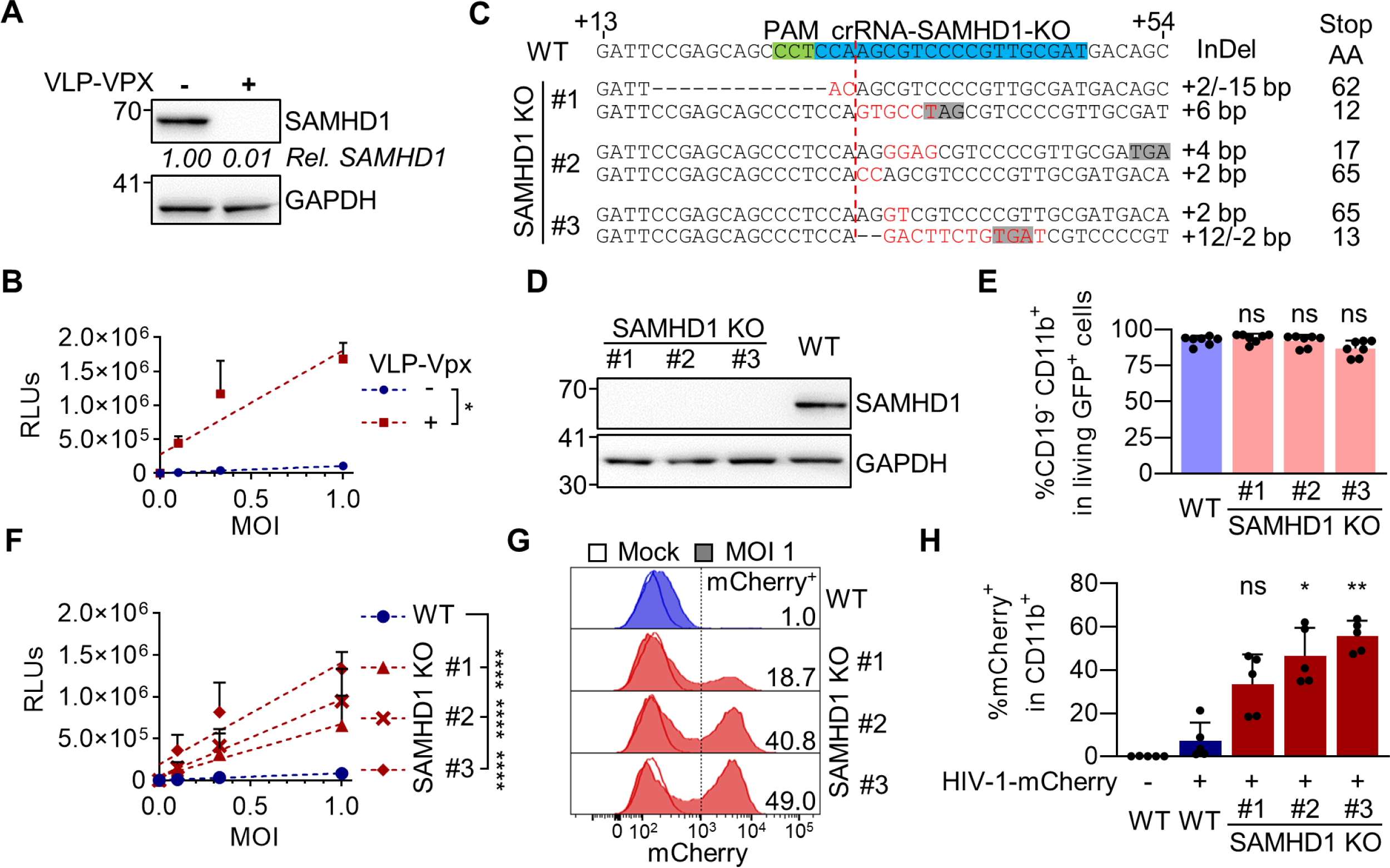
SAMHD1 restricts HIV-1 replication in transdifferentiated BLaER1 cells. **(A)** Transdifferentiated BLaER1 cells were treated with VLP-Vpx or medium for 24 h. SAMHD1 degradation was measured by immunoblot and quantified relative to GAPDH expression, followed by normalization to medium treated control (mean of n = 3). **(B)** VLP-Vpx or medium treated transdifferentiated BLaER1 cells were infected with VSV-G pseudotyped HIV-1 single-cycle luciferase reporter virus pNL4.3 E^-^ R^-^ luc at an MOI of 0.1, 0.33 and 1. Relative light units (RLUs) were quantified by luciferase measurement at 24 hpi. Linear regressions (dashed lines) were calculated and differences of slopes were tested for significance (n = 3, t-test). **(C)** BLaER1 cells were treated with CRISPR/Cas9 protein complexed with crRNA-SAMHD1-KO. Single cell clones were Sanger sequenced after TA-cloning to separate alleles and aligned to WT sequence. Insertions (red) and/or deletions (InDel) are indicated, as well as the position of the premature stop codon (gray), introduced by the respective genetic modification. **(D)** Genetically confirmed SAMHD1 knock-out (KO) clones were analyzed via immunoblot for SAMHD1 expression in transdifferentiated BLaER1 cells. GAPDH was used as loading control (n = 7). **(E)** Percentages of CD19^-^ CD11b^+^ cells in viableGFP^+^ transdifferentiated WT and SAMHD1 KO cells were quantified by flow cytometry (n = 7, One-way ANOVA). **(F)** RLUs in transdifferentiated BLaER1 WT and KO cell clones were quantified 24 hpi with pNL4.3 E^-^ R^-^ luc (VSV-G). Statistical significance of differences between linear regressions (dashed lines) in SAMHD1 KO clones compared to WT are indicated (n = 7, One-way ANOVA). **(G, H)** Transdifferentiated WT and SAMHD1 KO cell clones were infected with VSV-G pseudotyped HIV-1 single cycle mCherry reporter virus pNL4.3 IRES mcherry E^-^ R^+^ at MOI 1. Percentage of mCherry^+^ cells was quantified by flow cytometry in viable GFP^+^ CD11b^+^ BLaER1 cells 24 hpi. **(G)** Representative histograms are shown for mock and HIV-1-mCherry reporter virus infected cells. Percentage of mCherry^+^ in viable GFP^+^ CD11b^+^ BLaER1 cells is indicated. **(H)** Bar graphs indicate mean of experiments, dots individual biological replicates (n = 5, Kruskal–Wallis test). **(B, E, F, H)** Error bars correspond to standard deviation (* p < 0.05; ** p < 0.01; *** p < 0.001; **** p < 0.0001; ns, not significant).

### A pipeline to generate mutants of SAMHD1 by CRISPR/Cas9 mediated knock-in

So far, mutagenic analysis of SAMHD1 has been limited to model systems in which SAMHD1 is overexpressed by transient transfection or retroviral transduction. Overexpression of SAMHD1, especially in the context of phosphomimetic T592E or phosphoablative T592A mutation and their effect on viral restriction and intracellular dNTP levels, might affect functional readouts due to non-physiological expression levels, abnormal genomic context and altered post-translational regulation (Majer et al. 2019). To overcome this challenge, we decided to introduce SAMHD1 point mutations directly into the *SAMHD1* gene locus by CRISPR/Cas9 KI. Therefore, we developed a pipeline based on the introduction of RNPs and single-stranded DNA correction templates by electroporation, followed by an allele-specific PCR (KASP-genotyping assay screening) and rigorous validation by Sanger sequencing and quantitative genomic PCR to exclude large genomic deletions (qgPCR) (Weisheit et al. 2020) (Fig. 3A). We identified single cell clones, displaying homozygous introduction of T592A and T592E mutations into the *SAMHD1* locus of BLaER1 cells (Fig. 3B). Quantification of allele numbers of SAMHD1 exon 16 revealed that the majority of homozygous single cell T592A and T592E KI clones still contained two alleles of SAMHD1 exon16 (Fig. 3C). However, we could identify one out of 8 clones analyzed (Clone X), which showed loss of one allele in qgPCR, indicative of pseudo-homozygosity (Weisheit et al. 2020). In total, we were able to generate and validate two homozygous T592A, as well as three homozygous T592E BLaER1 KI mutants, corresponding to a homozygous KI frequency of ∼1% and highlighting the necessity of KASP-screening to reduce the number of KI candidates (Fig. 3D). Expression of SAMHD1 mutants in transdifferentiated T592A or T592E KI BLaER1 single cell clones was at similar level compared to WT protein in the respective parental cell line (Fig. 3E). SAMHD1 KI had no negative impact on BLaER1 transdifferentiation (Fig. 3F). In summary, using our pipeline we introduced homozygous T592A and T592E mutations into the endogenous *SAMHD1* locus of BLaER1 cells without affecting SAMHD1 expression or BLaER1 transdifferentiation into macrophage-like cells.

**Figure 3:**
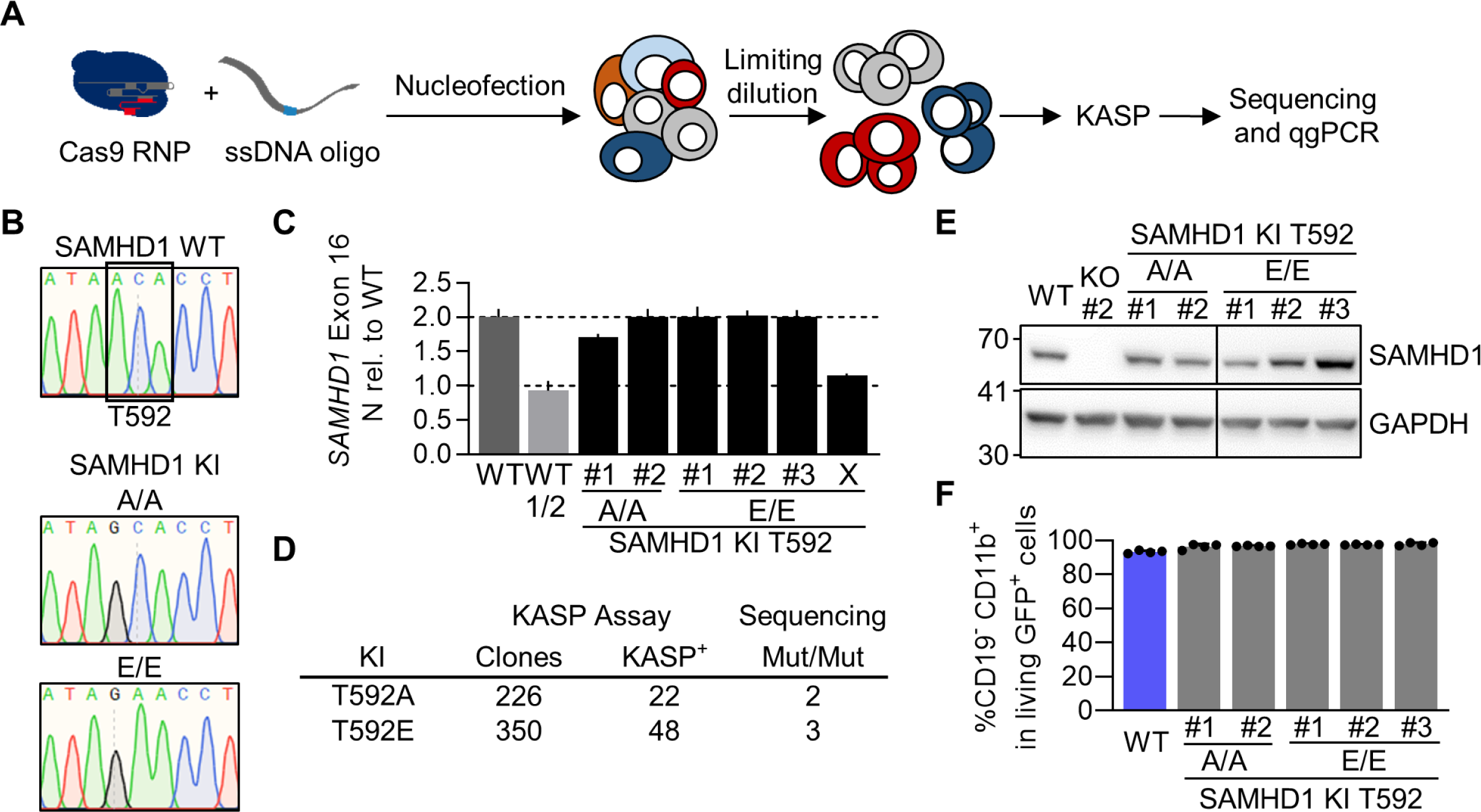
A pipeline to generate mutants of SAMHD1 by CRISPR/Cas9 mediated knock-in. **(A)** Schematic representation of CRISPR/Cas9 meditated knock-in (KI) to generate mutants of SAMHD1 in BLaER1 cells. Cas9 ribonucleoprotein (RNP) together with ssDNA correction template was introduced into BLaER1 cells via nucleofection. Single cell clones generated by limiting dilution were screened using KASP assay and validated by Sanger sequencing and quantitative genomic PCR (qgPCR) (Weisheit et al. 2020). **(B)** Representative sections of Sanger sequencing traces obtained from genomic *SAMHD1* exon 16, highlighting successful bi-allelic single base exchange at the base triplet corresponding to amino acid position T592 in BLaER1 SAMHD1 KI T592A and T592E mutant single cell clones. No further mismatches were detected up- or downstream of shown section in the amplified region. Two independent sequencing runs were performed. Homozygous T592E mutants were additionally confirmed by allele specific sequencing after TA-cloning. **(C)** Quantitative genomic PCR for *SAMHD1* exon 16 against reference gene *TERT* was performed and 2^-Δct^ value obtained from SAMHD1 KI clones normalized to WT in order to obtain the allele number. As a control half of the WT (WT 1/2) DNA was inoculated and Δct of *SAMHD1* was calculated against ct of *TERT*, which was obtained in the WT with normal DNA amount. Error bars indicate standard deviation of technical triplicates in a representative experiment (n = 2). **(D)** Number of single cell clones obtained from CRISPR/Cas9 RNP and ssDNA correction oligo treated BLaER1 cells and number of clones scoring positive in KASP assay, as well as homozygous (Mut/Mut) mutants identified by Sanger sequencing and confirmed by qgPCR are shown. **(E)** Transdifferentiated SAMHD1 KI BLaER1 cells were analyzed by immunoblot for SAMHD1 expression and compared to WT cells. GAPDH was used as a loading control (representative for n = 3). **(F)** Percentage of CD19^-^ CD11b^+^ cells in viable GFP^+^ transdifferentiated WT and SAMHD1 KI cells were quantified by flow cytometry. Error bars indicated standard deviation of biological replicates (n = 4).

### Homozygous SAMHD1 T592E mutation increases HIV-1 infection in transdifferentiated BLaER1 cells

We infected several clones of transdifferentiated homozygous SAMHD1 phosphoablative T592A and phosphomimetic T592E KI BLaER1 cell mutants with HIV-1-mCherry reporter virus and measured the fold change of %mCherry^+^ cells in CD19^+^ viable GFP^+^ cells relative to infection in WT cells. In all three clones, homozygous SAMHD1 T592E mutation significantly increased HIV-1-mCherry infection up to 31-fold (T592E/T592E Clone #1 *p = 0.0017*, #2 and #3 *p < 0.0001*, n = 3, One-way ANOVA) (Fig. 4A and 4B). In contrast, SAMHD1 T592A KI mutants completely retained their restrictive potential in transdifferentiated BLaER1 cell clones and behaved similar to WT BLaER1 cells upon challenge with HIV-1-mCherry (Fig. 4A and 4B). Using CRISPR/Cas9 KI, we were able to validate the loss of HIV-1 restriction in SAMHD1 phosphomimetic T592E mutants in macrophage-like cells. In this model, mutants of SAMHD1 are analyzed in the native genomic context and show physiological expression levels, confirming the role of T592 phosphorylation in the regulation of the anti-viral activity of SAMHD1.

**Figure 4:**
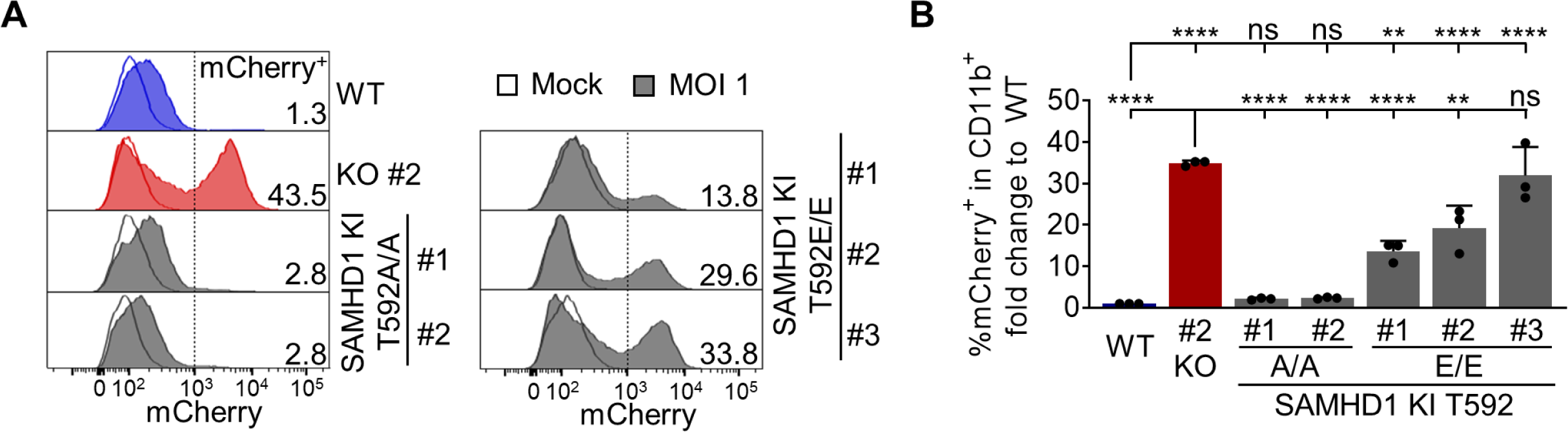
Homozygous SAMHD1 T592E, but not T592A mutation leads to loss of HIV-1 restriction in transdifferentiated BlaER1 cells. **(A, B)** Transdifferentiated homozygous SAMHD1 T592E and T592A BLaER1 KI clones were infected with VSV-G pseudotyped HIV-1 single cycle mCherry reporter virus pNL4.3 IRES mcherry E^-^ R^+^ at MOI 1. Percentage of mCherry^+^ cells was quantified by flow cytometry in viable GFP^+^ CD11b^+^ BLaER1 cells at 24 hpi. **(A)** Representative histograms are shown for mock and HIV-1-mCherry reporter virus infected cells. Percentage of mCherry^+^ cells in viable GFP^+^ CD11b^+^ BLaER1 cells is indicated (n = 3). **(B)** To calculate fold change, percentage of mCherry^+^ cells in infected SAMHD1 KI clones was normalized to WT. Bar graphs indicate mean of experiments, dots individual biological replicates. Error bars correspond to standard deviation (n = 3, One-way ANOVA, ** p < 0.01; **** p < 0.0001; ns, not significant).

### SAMHD1 T592E or T592A knock-in does not affect dNTP levels in transdifferentiated BLaER1 cells

Previous reports on the effect of SAMHD1 T592 phosphorylation on SAMHD1 dNTPase activity were in-conclusive (Majer et al. 2019). In order to correlate HIV-1 restrictive potential in transdifferentiated BLaER1 cells with cellular dNTP pool size and thus SAMHD1 dNTPase activity, we measured intracellular dNTP levels by primer extension assay. Transdifferentiated WT BLaER1 cells contained low amounts of dATP (846 ± 63 fmol/10^6^ cells, n = 5), dCTP (788 ± 117 fmol/10^6^ cells, n = 5), dGTP (724 ± 94 fmol/10^6^ cells, n = 5) and dTTP (933 ± 342 fmol/10^6^ cells, n = 5). Depletion of the minor fraction of CD19^+^ cells after transdifferentiation further reduced the levels of dATP (578 fmol/10^6^ cells), dCTP (661 fmol/10^6^ cells), dGTP (295 fmol/10^6^ cells) and dTTP (448 fmol/10^6^ cells). Since activity of HIV-1 RT is likely to be dependent on cellular dNTP concentrations rather than total dNTP pools, we determined cellular dNTP concentrations as a function of transdifferentiated BLaER1 cell volumes (569 ± 138 µm^3^, n_cells_ = 15). We found transdifferentiated WT BLaER1 cells to harbor dNTP concentrations (Tab. 1), similar or lower to those found in resting T cells (Diamond et al. 2004). Depletion of incompletely transdifferentiated (CD19^+^) cells from bulk preparations of transdifferentiated BLaER1 cells further reduced dNTP concentrations (Tab. 1). As expected, SAMHD1 KO led to a significant increase in cellular dATP (2.3-fold, *p < 0.0001*, One-way ANOVA), dGTP (3.2-fold, *p < 0.0001*, One-way ANOVA) and dTTP (2.2-fold, *p < 0.0001*, One-way ANOVA) levels in transdifferentiated BLaER1 cells, as compared to WT cells (Fig. 5A). In contrast, neither homozygous SAMHD1 T592E, nor T592A mutations led to an increase of cellular dNTP levels (Fig. 5A). Since SAMHD1 KO only slightly affected cellular dCTP levels, dNTP composition in transdifferentiated BLaER1 SAMHD1 KO cells was altered. In contrast, neither SAMHD1 T592E nor SAMHD1 T592A KI mutants showed consistent differences in cellular dNTP composition (Fig. 5B). In summary, dNTP measurements in transdifferentiated BLaER1 cells, harboring homozygous phosphomimetic T592E or phosphoablative T592A mutations in the endogenous *SAMHD1* locus indicate that phosphorylation at SAMHD1 residue T592 has no impact on cellular dNTP pools and is therefore unlikely to regulate SAMHD1 dNTPase activity in cells.

**Figure 5:**
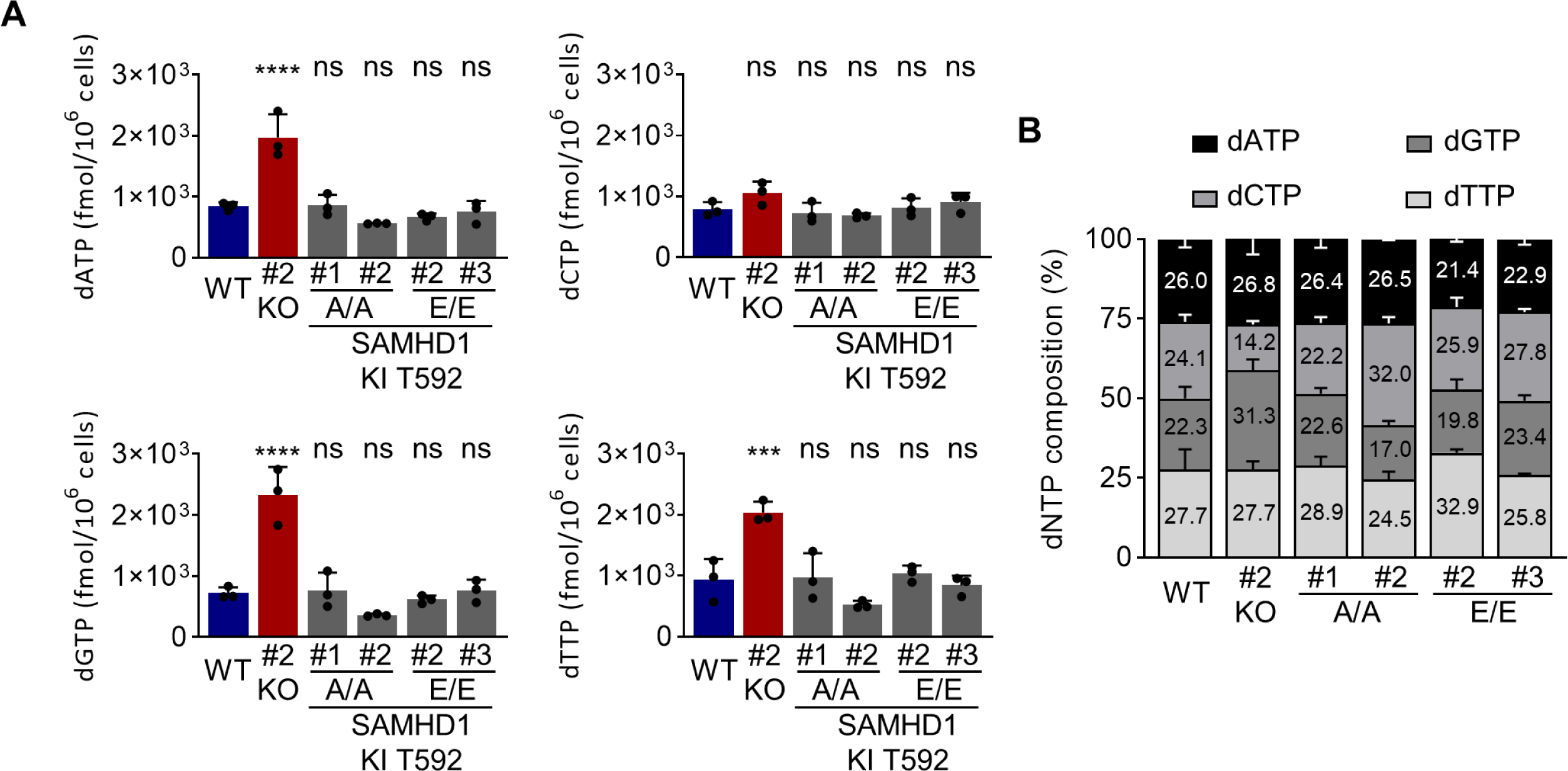
SAMHD1 T592E or T592A knock-in does not affect dNTP levels in transdifferentiated BLaER1 cells. **(A, B)** Cellular dNTP levels were measured in transdifferentiated homozygous SAMHD1 T592E and T592A BLaER1 KI mutants. dNTP amounts were compared to transdifferentiated WT BLaER1 cells. **(A)** Amount of indicated dNTP is depicted per 1 x 10^6^ cells. Bar graphs indicate mean of experiments, dots individual biological replicates. Error bars correspond to standard deviation (n = 3, One-way ANOVA, *** p < 0.001; **** p < 0.0001; ns, not significant). **(B)** dNTP composition in individual BLaER1 SAMHD1 KI clones is shown, with total dNTP content set as 100%. Error bars indicate standard deviation (n = 3).

**Table 1:**
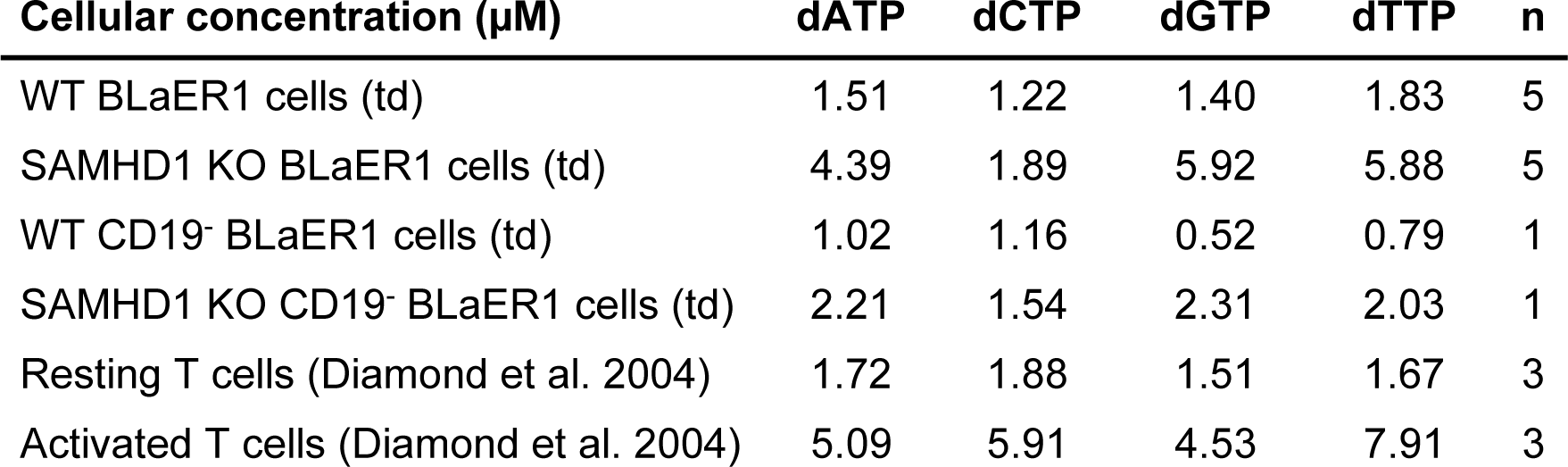
dNTP concentrations in transdifferentiated and CD19 depleted BLaER1 cells.

### SAMHD1 T592E knock-in relieves a block to HIV-1 reverse transcription and HIV-1 infection, which is not further enhanced by SAMHD1 T592E mutant depletion

Previous results indicate that SAMHD1 inhibits HIV-1 replication at the step of reverse transcription (Hrecka et al. 2011; Schott et al. 2018; Baldauf et al. 2017). To better understand the effect of endogenous SAMHD1 T592E mutation on HIV-1 replication, we measured the abundancy of HIV-1-mCherry reverse transcription products in CD19-depleted transdifferentiated BLaER1 cells. While late reverse transcription product copy numbers in WT BLaER1 cells stayed low until 24 hpi, both SAMHD1 KO and SAMHD1 T592E KI mutant BLaER1 cells showed a strong increase in HIV-1 late reverse transcription product copy numbers starting from 9 hpi (Fig. 6A). This indicates that endogenous mutation of SAMHD1 T592E in transdifferentiated BLaER1 cells relieves a block to HIV-1 replication which is situated at or before the step of reverse transcription. Several restriction mechanisms have been proposed for SAMHD1, notably dNTP degradation, RNAse activity and nucleic acid binding (Choi et al. 2015; Ryoo et al. 2014; Beloglazova et al. 2013; Goncalves et al. 2012; Seamon et al. 2015; Goldstone et al. 2011; Powell et al. 2011; Franzolin et al. 2013; Majer et al. 2019). To understand if additional SAMHD1 mediated, but T592 de-phosphorylation independent anti-lentiviral mechanisms could influence HIV-1 replication in transdifferentiated BLaER1 cells, we tested whether SAMHD1 mutant depletion using VLP-Vpx would further enhance HIV-1 replication. As anticipated, VLP-Vpx mediated SAMHD1 depletion (Fig. 6B) in WT cells significantly increased the percentage of HIV-1-mCherry infected CD11b^+^ macrophage-like BLaER1 cells, while there was no effect on SAMHD1 KO cells, when compared to empty VLP treated control cells (Fig. 6C; WT *p < 0.0001*, KO Clone #2 *p = 0.9615*, n = 3, One-way ANOVA). In contrast, depletion of endogenous T592A mutant SAMHD1 significantly increased HIV-1 infection rates (Clone #1 and 2 *p < 0.0001*, n = 3, One-way ANOVA). VLP-Vpx treatment in SAMHD1 T592E KI mutant clones however had no effect on HIV-1 infection rates (Fig. 6C; Clone #2 *p = 0.2654*, #3 *p = 0.1788*, n = 3, One-way ANOVA). Thus, our data indicate that no T592 de-phosphorylation independent anti-lentiviral mechanism is exerted by SAMHD1 in macrophage-like cells.

**Figure 6:**
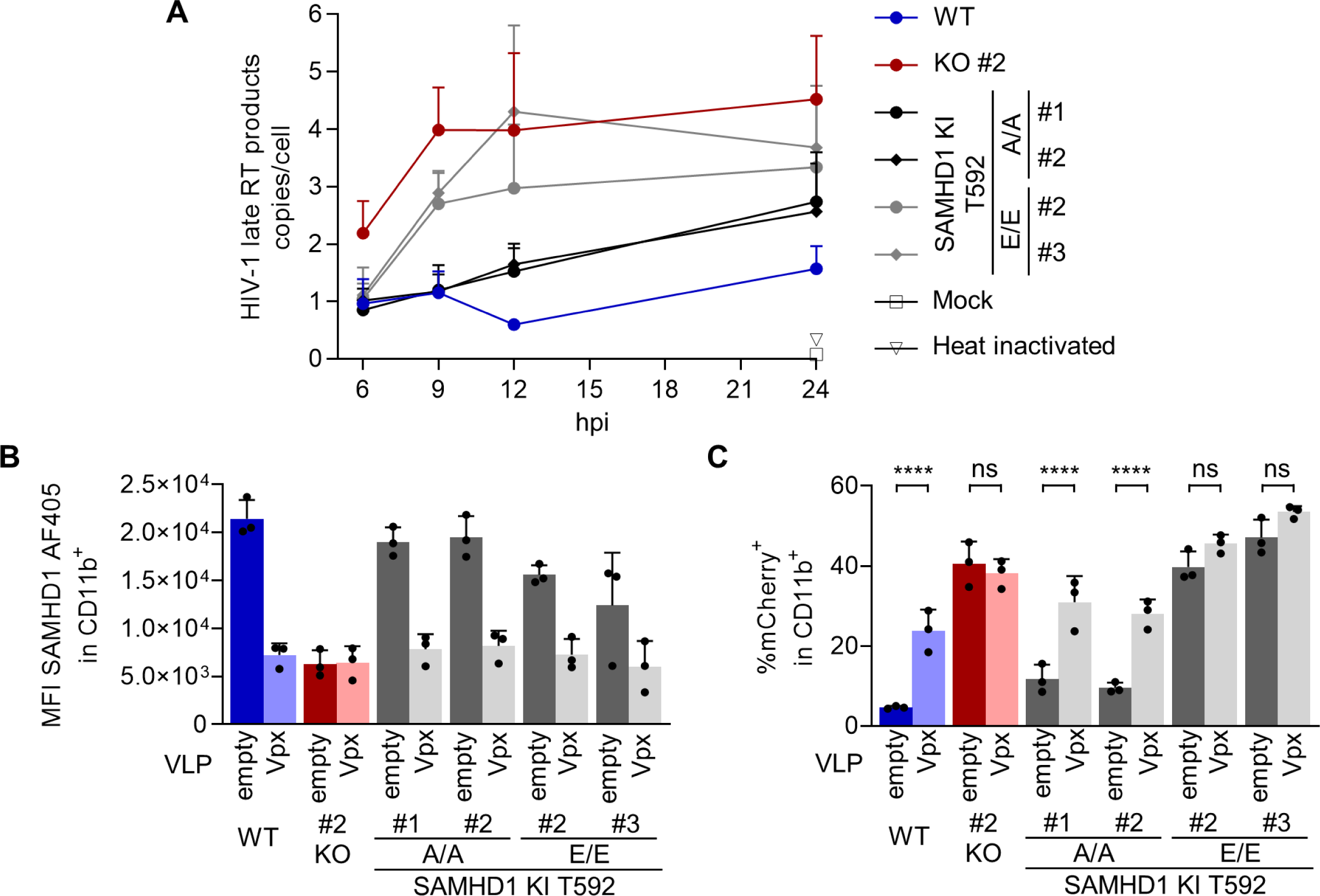
SAMHD1 T592E knock-in relieves a block to HIV-1 reverse transcription and HIV-1 infection, which is not further enhanced by SAMHD1 T592E mutant depletion. **(A)** Transdifferentiated homozygous SAMHD1 T592E and T592A BLaER1 KI clones were depleted for CD19^+^ cells and infected with VSV-G pseudotyped HIV-1 single cycle mCherry reporter virus pBR HIV1 M NL4.3 IRES mcherry E^-^ R^+^ at MOI 1. At the indicated time point post infection, late reverse transcription (RT) products were quantified by qPCR and normalized against *PBGD* to determine late RT copy number per cell. Error bars correspond to standard deviation (n = 3). **(B, C)** Transdifferentiated homozygous SAMHD1 T592E and T592A BLaER1 KI clones were treated with virus-like particles (VLPs) co-packaging Vpx or empty controls in parallel to infection with HIV-1-mCherry (MOI 1). **(B)** SAMHD1 abundancy in CD11b^+^ cells was analyzed by flow cytometry at 24 hpi. **(C)** Percentage of mCherry^+^ cells in CD11b^+^ BLaER1 cells at 24 hpi is shown. Bar graphs indicate mean of experiments, dots individual biological replicates. Error bars correspond to standard deviation (n = 3, One-way ANOVA; **** p < 0.0001; ns, not significant).

### The integrity of the catalytic site in SAMHD1 is critical for anti-viral activity, however antiviral activity correlates poorly with global cellular dNTP levels

Previous results indicate the loss of SAMHD1’s anti-viral potential in mutants of key catalytic residues, as shown by overexpression of mutants such as combined H206A_D207A or the D311A mutant (Laguette et al. 2011; Lahouassa et al. 2012; Arnold et al. 2015; White et al. 2013a). However, overexpression might introduce substantial experimental bias, therefore, we decided to validate the requirement of the catalytic dNTPase pocket by CRISPR/Cas9 knock-in. To do so, we modified the residues, H210, D218 and D311 in BLaER1 cells. These residues have been suggested to be directly or indirectly implicated in the triphosphohydrolase activity of SAMHD1 (Morris et al. 2020). The introduction of homozygous mutations were verified by Sanger sequencing and qgPCR (Fig. S1). The mutant proteins with alanine substitution of the respective residues, were expressed in macrophage-like BLaER1 cells (Fig. 7A). While H210A showed similar or slightly higher expression compared to WT protein, D218A and D311A expression was slightly reduced. Importantly, we did not see increased SAMHD1 pT592 in any of the mutant clones, indicating phospho-regulation of all three endogenous catalytic site mutants similar to WT (Fig. 7A). As expected, macrophage-like BLaER1 cells harboring SAMHD1 D311A mutation displayed a significant increase in cellular dNTP levels (dATP #1-4 *p<0.0001*; dCTP #1 *p=0.0079*, #2 *p=0.0003*, #3 *p=0.1271*, #4 *p=0.0169*; dGTP #1-4 *p<0.0001*; dTTP #1-4 *p<0.0001*, n = 3, One-way ANOVA) (Fig. 7 B to E), for dATP, dGTP and dTTP even above the levels measured in SAMHD1 KO BLaER1 cell clone #2, indicating a complete loss of SAMHD1 dNTPase activity in D311A mutant protein. The increase of dNTP levels in homozygous D210A and D218A mutant cells was much less pronounced and only consistently significant in D210A clone #2 and D218A clone #1 for dATP, dGTP and dTTP (dATP D210A #2 *p=0.0025*, D218A #1 *p=0.0724;* dGTP D210A #2 *p<0.0001*, D218A #1 *p=0.0003;* dTTP D210A #2 *p=0.0052*, D218A #1 *p=0.0457*, n = 3, One-way ANOVA) with levels similar to SAMHD1 KO (Fig. 7 B to C). Importantly, several cell clones, namely D210A #3 and 4, as well as D218A #2, revealed dNTP levels similar or only marginally increased, when compared to WT or T592A mutant cells indicating that D210A and D218A mutation does not lead to complete loss of SAMHD1 dNTPase activity or alternatively, loss of control of dNTP levels could be compensated. Next, we infected transdifferentiated D210A, D218A and D311A mutant cells with HIV-1-mCherry reporter virus at three different MOI and quantified infected mCherry^+^ cells in CD11b^+^ BLaER1 cells after 24 h (Fig. 8). CRISPR/Cas9 D210A and D311A KI mutant cells showed a significant (*p<0.0001* for all clones, n = 3, One-way ANOVA) increase in infection level, which was consistent over a range of MOI from 0.1 to 1 (Fig 8B). Surprisingly, however, the loss of restrictive potential in the D218A mutant cell clones was strikingly lower and only significant for Clone #2 (#1 *p=0.1975*, #2 *p<0.0001*, n = 3, One-way ANOVA). When compared to SAMHD1 KO, HIV-1-mCherry infection at MOI 1 was significantly lower in D218A (*p<0.0001*, n = 3, One-way ANOVA), but similar in D210A and D311A mutants. Depletion of SAMHD1 D210A and D311A mutant protein by VLP-Vpx treatment did not further increase HIV-1-mCherry infection at MOI 0.1 (Fig 8C and D). In contrast, VLP-Vpx mediated depletion of SAMHD1 D218A mutants allowed higher infection levels, which after depletion were similar to SAMHD1 KO cells (Fig. 8D). In summary, using CRISPR/Cas9 KI mutants of the SAMHD1 catalytic dNTP pocket, we were able to show that the residues H210 and D311 are critical for anti-viral restriction. However, this seems not to be the case for D218, highlighting that the individual residues of the catalytic pocket might be differentially involved in HIV-1 restriction. Importantly, lack of correlation of cellular dNTP levels and HIV-1 restriction potential in D210A and D218A mutants, together with loss of HIV-1 restriction in dNTP low T592E mutants, indicates that maintenance of low global dNTP levels is a poor predictor of SAMHD1 anti-viral activity.

**Figure 7:**
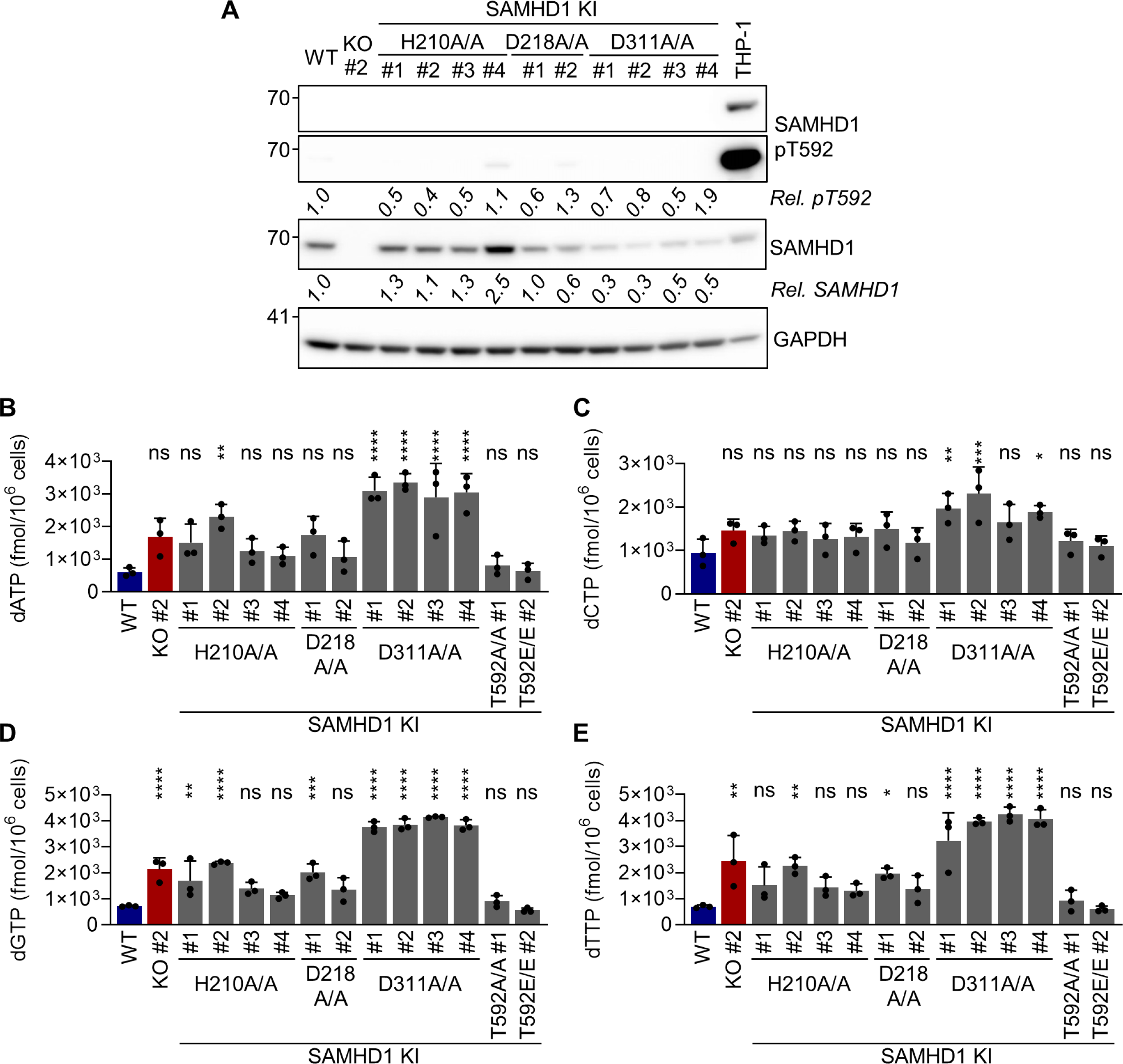
CRISPR/Cas9 mediated mutation of SAMHD1 catalytic core residues partly increases dNTP levels in transdifferentiated BLaER1 cells. **(A)** Transdifferentiated SAMHD1 KI BLaER1 cells were analyzed by immunoblot for SAMHD1 expression and T592 phosphorylation in comparison to WT and cycling THP-1 cells. GAPDH was used as a loading control. Mean of SAMHD1 T592 phosphorylation (pT592) relative to total SAMHD1 expression and SAMHD1 expression relative to GAPDH abundancy in transdifferentiated BLaER1 cells was normalized to WT cells (n = 4). **(B-E)** Cellular dNTP levels were measured in transdifferentiated homozygous SAMHD1 H210A, D218A and D311A BLaER1 KI mutants. dNTP amounts were compared to transdifferentiated WT and SAMHD1 KO BLaER1 cells. Amount of indicated dNTP is depicted per 1 x 10^6^ cells. Bar graphs indicate mean of experiments, dots individual biological replicates. Error bars correspond to standard deviation (n = 3, One-way ANOVA, * = p < 0.05; ** = p <0.01; *** p < 0.001; **** p < 0.0001; ns, not significant).

**Figure 8:**
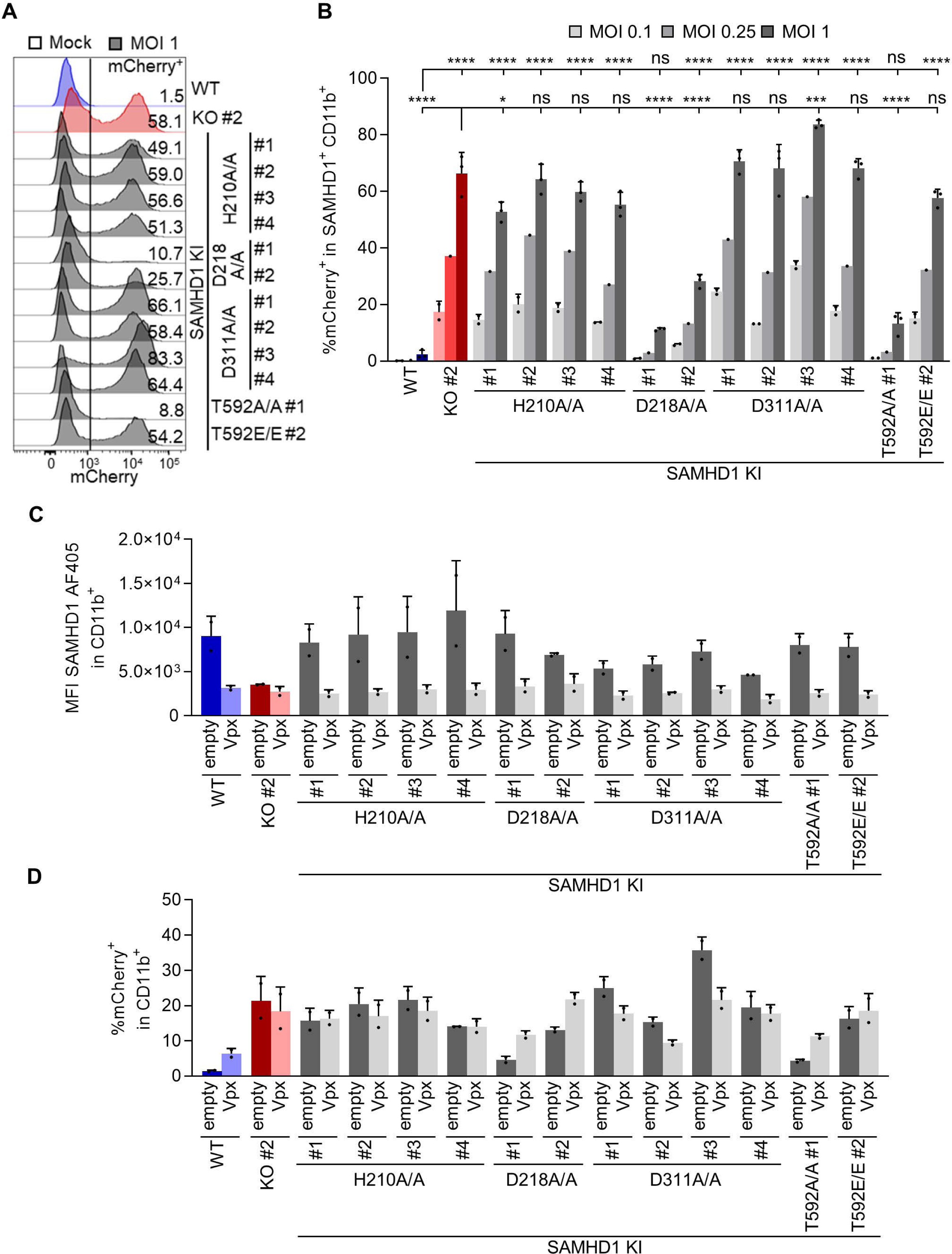
CRISPR/Cas9 mediated mutation of SAMHD1 catalytic core residues reveal that the integrity of the catalytic site is required for HIV-1 restriction. **(A, B)** Transdifferentiated homozygous SAMHD1 T592A, T592E, H210A, D218A and D311A BLaER1 KI clones were infected with VSV-G pseudotyped HIV-1 single cycle mCherry reporter virus pBR HIV1 M NL4.3 IRES mcherry E^-^ R^+^ at MOI 1, 0.25 and 0.1, as indicated. Percentage of mCherry^+^ cells was quantified by flow cytometry in CD11b^+^ SAMHD1^+^ BLaER1 cells at 24 hpi. **(A)** Representative histograms are shown for HIV-1-mCherry reporter virus infected cells. Percentage of mCherry^+^ cells in CD11b^+^ SAMHD1^+^ BLaER1 cells is indicated (n = 3). **(B)** Percentage of mCherry^+^ cells in infected SAMHD1 KI clones is shown for MOI 0.1 (n = 2), 0.25 (n = 1) and 1 (n = 3). **(C, D)** Transdifferentiated homozygous SAMHD1 T592A, T592E, H210A, D218A and D311A BLaER1 KI clones were treated with virus-like particles (VLPs) co-packaging Vpx or empty controls in parallel to infection with HIV-1-mCherry (MOI 0.1). **(C)** SAMHD1 abundancy in CD11b^+^ cells was analyzed by flow cytometry at 24 hpi. **(D)** Percentage of mCherry^+^ cells in CD11b^+^ BLaER1 cells at 24 hpi is shown. Bar graphs indicate mean of experiments, dots individual biological replicates. Error bars correspond to standard deviation (n = 2). Bar graphs indicate mean of experiments, dots individual biological replicates. Error bars correspond to standard deviation (One-way ANOVA, * p < 0.01; *** p < 0.001; **** p < 0.0001; ns, not significant).

## Discussion

SAMHD1 is a major cellular dNTPase and a potent HIV-1 restriction factor (Laguette et al. 2011; Berger et al. 2011; Hrecka et al. 2011; Baldauf et al. 2012; Descours et al. 2012; Goldstone et al. 2011; Franzolin et al. 2013). However, whether SAMHD1 dNTPase activity is meditating its anti-viral activity and how this is regulated by T592 phosphorylation is still a matter of debate (Majer et al. 2019; Welbourn und Strebel 2016).

To tackle this question with a novel toolkit, we used transdifferentiated BLaER1 cells as an alternative and versatile myeloid model to study HIV-1 infection in macrophage-like cells. Transdifferentiated BLaER1 cells closely resemble human macrophages and have successfully been used to study innate immune signaling (Gaidt et al. 2016; Rapino et al. 2013). In contrast to PMA activated THP-1 or U937 cells, BLaER1 cell transdifferentiation relies on the activation of the fusion protein of the macrophage transcription factor C/EBPα and the estradiol receptor leading to the induction of a myeloid cell program (Rapino et al. 2013; Gaidt et al. 2018). Transdifferentiated BLaER1 cells show surface expression of classical monocyte-derived macrophage and dendritic cell markers CD14, CD163, CD206 and CD11c, validating previous transcriptional data (Rapino et al. 2013), as well as HIV-1 CD4, CCR5 and CXCR4 (co-) receptor expression. This cell system is an interesting physiological model for HIV-1 infection in macrophage-like cells, as we were able to show that SAMHD1 is completely dephosphorylated at residue T592 in transdifferentiated BLaER1 cells and serves as a major restriction factor for HIV-1.

To test mutants of SAMHD1 for anti-viral restriction activity in a physiological genetic and relevant cellular context, we combined transdifferentiated BLaER1 cells, as a novel myeloid model, with CRISPR/Cas9 mediated knock-in. We developed a gene editing strategy using CRISPR/Cas9 RNP and ssDNA oligo introduction by nucleofection, KASP screening of single cell clones and rigorous validation by sequencing and qgPCR (Fig. 3A). By introducing point mutations that correspond to phosphoablative T592A and phosphomimetic T592E mutations into the genomic *SAMHD1* locus, we were able to genetically uncouple SAMHD1 mediated anti-viral restriction and cellular dNTPase activity. As expected, we showed that endogenous T592 phospho-mimetic mutants lose their anti-viral activity against HIV-1 in macrophage-like BLaER1 cells, while phospho-ablative KI mutants maintained their anti-viral potential (Fig. 4). Furthermore, using endogenous mutants of T592, we highlighted that T592 phospho-regulation indeed affects anti-lentiviral activity of SAMHD1 at the step of reverse transcription (Fig. 6A). Thus, the first time, we both validated and expanded our knowledge on the phenotypic consequences of SAMHD1 T592E phosphomimetic mutation, and hence the effect of T592 phosphorylation, for its anti-viral restriction activity in a myeloid model not based on overexpression.

Overexpression of SAMHD1 mutants in U937 background using retroviral transduction has several technical limitations. Even though PMA-activated U937 cells are often considered not to express SAMHD1, they actually can express small amounts of endogenous SAMHD1, which can be further enhanced upon interferon treatment (Riess et al. 2017). Presence of endogenous WT SAMHD1 might affect the function of overexpressed mutant SAMHD1, especially if heterotetramers are formed. In addition, strong exogenous, often viral promotors drive SAMHD1 expression here, leading to high expression levels and to non-physiological phosphorylation ratios, *i.e.* hyperphosphorylation (data not shown). In mutants generated by CRISPR/Cas9 KI, modified SAMHD1 is expressed in the physiological genomic context from the endogenous promotor and thus under normal transcriptional regulation. In BLaER1 cells, SAMHD1 expression is very low in native cells, but strongly induced upon transdifferentiation (Fig. 1D). This is also the case for SAMHD1 mutants. The use of CRISPR/Cas9 KIs avoids potential effects of constitutively expressed SAMHD1 mutants on cycling BLaER1 cells.

In macrophage-like WT BLaER1 cells, we measured dNTP levels and concentrations, which were similar or slightly lower than those found in resting T cells (Tab. 1) (Diamond et al. 2004). After depletion of CD19^+^ incompletely transdifferentiated cells from bulk preparations of transdifferentiated BLaER1 cells, we were able to further reduce the levels of all dNTPs (Tab. 1) (Diamond et al. 2004). Considering HIV-1 reverse transcriptase K_m_ and K_d_ values measured *in vitro*, this indicates that the dNTP concentrations found in transdifferentiated BLaER1 cells are sufficiently low to restrict or delay HIV-1 RT (Kennedy et al. 2010; Klarmann et al. 1993; Derebail und DeStefano 2004; Jacques et al. 2016).

Concomitantly, SAMHD1 KO increased cellular dNTP concentrations in transdifferentiated BLaER1 cells up to 4-fold (Fig. 5A), which is reminiscent of the 5- to 8-fold increase upon T cell activation (Diamond et al. 2004). In stark contrast however, neither endogenous SAMHD1 T592E nor T592A mutation increased cellular dNTP concentrations in transdifferentiated BLaER1 cells (Fig. 5A). This indicates that the loss of restriction observed in endogenous T592E mutants is probably not caused by increased dNTP levels or reduced SAMHD1 dNTPase activity in transdifferentiated BLaER1 cells. This lack of correlation between cellular dNTP levels and HIV-1 restriction was also observed in a study published during revision, highlighting the possibility that T592 phosphorylation might indeed impact SAMHD1 tetramer stability (Monit et al. 2019). However, using our endogenous approach, we exclude artifacts of overexpression and the use of U937 cells as a model and thereby improve the physiological value of the conclusions.

In addition, we demonstrated that SAMHD1 T592E and T592A mutations had no consistent effect on dNTP pool composition in macrophage-like BLaER1 cells (Fig. 5B), ruling out an effect of the phosphomimetic mutation on SAMHD1 dNTPase substrate preferences and thus dNTP ratios, which was proposed earlier (Jang et al. 2016). More specifically, endogenous SAMHD1 T592E mutations did not increase cellular dCTP concentration (Fig. 5A). Taken together mutagenic analysis of SAMHD1 residue T592 indicates that SAMHD1 dNTPase activity or substrate preference in transdifferentiated BLaER1 cells is not regulated by phosphorylation at this specific residue. Consequently, loss of HIV-1 restriction in SAMHD1 T592E mutants cannot be attributed to changes in SAMHD1 dNTPase activity. In addition, by combining SAMHD1 T592E KI mutants with VLP-Vpx mediated depletion of SAMHD1 in trans, we did not find an anti-lentiviral activity of SAMHD1 independent of SAMHD1 T592 de-phosphorylation (Fig. 6B and C).

To address the relationship between SAMHD1 enzymatic function and its anti-lentiviral activity further, we introduced KI mutations for specific residues in the catalytic dNTPase pocket of SAMHD1 (Fig. 7A) (Morris et al. 2020). Loss of restrictive potential of H210A and D311A mutations in an endogenous context, indicate that the integrity of the catalytic site is indeed required for HIV-1 restriction (Fig. 8A and B). While the loss of restriction with the endogenous H210A and D311A mutations is in concordance with previous data employing overexpression models (Lahouassa et al. 2012; Laguette et al. 2011; Arnold et al. 2015; White et al. 2013a), two alternative hypotheses can be drawn from this observation: either the enzymatic SAMHD1 dNTPase activity per se is required for the inhibition of HIV-1 replication, or an (enzymatic) activity other than the canonical dNTPase activity, but still involving these residues is at play.

In favor of the first and so far, most often discussed hypothesis speaks data from further studies, showing that addition of exogenous dNs and a concomitant increase in cellular dNTP levels can leverage SAMHD1 mediated block to HIV-1 restriction (Baldauf et al. 2012; Lahouassa et al. 2012). Also, in concordance with previous results, the D311A mutation increases cellular dNTPs to levels equal or higher than those found in SAMHD1 KO macrophage-like BLaER1 cells (Fig. 7B to E), correlating HIV-1 restriction to SAMHD1 dNTPase activity. In addition, no D311 residue independent HIV-1 restriction potential was observed (Fig. 8C and D).

In contrast however, data on H210A and D218A mutants seems to be at odds with this hypothesis. First, while all SAMHD1 H210A mutant clones showed complete loss of HIV-1 restriction potential (Fig. 8 B and D), enhancement of dNTP levels was less pronounced (Fig. 7B to E). In particular, clone #3 and 4 showed global dNTP levels comparable to WT or T592A mutant clones, indicating that the H210 residue is not essential for dNTP degradation and that loss of dNTPase activity can only be partial in H210A mutant cells. Even more striking is the phenotype of the D218A mutation. SAMHD1 D218 was recently shown to be involved in the triphosphohydrolase reaction (Morris et al. 2020). D218A mutation only leads to partial loss of dNTPase activity in vitro (Morris et al. 2020), which is in concordance with our data in cellulo, showing a clone dependent significant increase of cellular dNTP levels, similar to what was found in H210A mutants (Fig. 7B to E). Importantly however, at least one of the D218A mutant cells maintained their potential to restrict HIV-1 replication (Fig. 8B and D). Thus, in the case of both H210A and D218A mutant cells, cellular dNTP levels and thus presumably SAMHD1 dNTPase activity, do not entirely correlate with their anti-lentiviral capacity.

A possible explanation is that a still to be defined function, other than dNTP triphosphohydrolase, which also depends on the residues of the catalytic pocket, is required for HIV-1 restriction and that this function is directly or indirectly modulated by T592 phosphorylation. Ribonuclease (RNase) activity was proposed as a mechanism responsible for HIV-1 restriction (Beloglazova et al. 2013; Ryoo et al. 2014; Choi et al. 2015). While initial reports of SAMHD1 associated RNase activity were strongly questioned by the community because of the reversible nature of SAMHD1 restriction and due to potential co-purification of unknown RNase(s) (Hofmann et al. 2013; Seamon et al. 2015; Antonucci et al. 2016; Tsai et al. 2023), recent evidence links SAMHD1 RNase activity to the inflammatory phenotype in Aicardi-Goutières syndrome (Maharana et al. 2022). Catalytic residues, such as H206, D207, D311 and H167 were proposed to be critical for RNase activity (Maharana et al. 2022; Beloglazova et al. 2013; Ryoo et al. 2014), while reports on the role of T592 phospho-regulation are contradictory (Ryoo et al. 2014; Maharana et al. 2022). The contribution of the proposed SAMHD1 RNase activity to HIV-1 restriction thus merits further investigation. Nucleic acid binding and exo-/endonuclease recruitment could be an alternative anti-lentiviral restriction mechanism. SAMHD1 is participating in the resolution of stalled replication forks and homologous recombination by recruitment of endo-/exonuclease MRE11 and CtIP (Coquel et al. 2018; Daddacha et al. 2017). Interestingly, SAMHD1 T592 phosphorylation is required for DNA end resection and resolution of stalled replication forks (Coquel et al. 2018). However, crucial residues of the catalytic dNTPase pocket such as H206, D207 and K312 of SAMHD1 seem dispensable for this process and endo-/exonuclease recruitment therefore unlikely to contribute to the phenotypes we observed in BLaER1 cells (Coquel et al. 2018; Daddacha et al. 2017). SAMHD1 was shown to bind to single-stranded RNA and DNA, as well as RNA or DNA with complex secondary structures in vitro (Beloglazova et al. 2013; Seamon et al. 2015). Whether nucleic acid biding is contributing to the recruitment of endo-/exonucleases in cells or anti-viral activity, is still an open question. Short phosphorothioate oligonucleotides can bind to the allosteric sites of SAMHD1, promoting formation of a distinct SAMHD1 tetramer (Yu et al. 2021). Mutations that abolish phosphorothioate oligonucleotide binding also reduced HIV-1 restriction potential. How SAMHD1 nucleic acid binding activity is regulated is currently not clear. SAMHD1 phosphomimetic T592E mutant had no influence on ssRNA and ssDNA binding in vitro (Seamon et al. 2015). Intriguingly, ssDNA or ssRNA binding to SAMHD1’s dimer-dimer interface can inhibit the formation of catalytically active SAMHD1 tetramer and thus interferes with dNTPase activity (Seamon et al. 2016).

Even though the SAMHD1-nucleic acid binding interface seems to be distinct from the catalytic pocket (Seamon et al. 2015; Yu et al. 2021), a deeper investigation is needed to understand this phenomenon and its implications for HIV-1 restriction in cells.

Another possible explanation could be that the dNTPase activity per se is required for HIV-1 restriction, but global cellular dNTP levels do not correlate with this anti-viral activity. Sub-cellular dNTP levels, i.e. in the nucleus or at sites of HIV replication, such as the nuclear pore, nuclear speckles or the capsid shell (Francis et al. 2020; Rensen et al. 2021) could be modulated by the active site mutants, such as H210A and D311A. In reverse, T592 phosphorylation could regulate the sub-cellular distribution of SAMHD1. T592 phosphorylation, however, has no influence on cytoplasmic vs. nuclear localization of SAMHD1 (White et al. 2013b). In addition, mutation of the n-terminal nuclear-localization signal (NLS) does not affect HIV-1 restriction (Brandariz-Nuñez et al. 2012). Yet, it is possible that T592 de-phosphorylation alters SAMHD1 distribution and i.e. allows the recruitment of SAMHD1 to the capsid shell or into the sub-nuclear compartments in which HIV-1 reverse transcription occurs and thereby locally modulates the availability of dNTPs.

SAMHD1 regulation certainly is more complex than commonly assumed. In addition to multiple potential phosphorylation sites, SAMHD1 is modified by acetylation, SUMOylation, ubiquitination and O-GlcNAcylation (White et al. 2013b; Welbourn et al. 2013; Kim et al. 2019; Ochoa et al. 2020; Lee et al. 2017; Lamoliatte et al. 2014; Hendriks et al. 2017; Lumpkin et al. 2017; Elia et al. 2015; Hu et al. 2021) and harbors redox-active cysteines (Mauney et al. 2017; Wang et al. 2018). Recently, SAMHD1 SUMOylation at residue K595 was shown to be required for HIV-1 restriction in PMA differentiated U937 cells. Overexpression of SAMHD1 mutants that abrogate restriction and SUMOylation at residue K595 did not show increased dATP levels, phenocopying phosphomimetic T592 mutants of SAMHD1 (Martinat et al. 2021). It will be interesting to investigate in more detail how T592 phosphorylation and K595 SUMOylation are integrated and it will be crucial to validate SAMHD1 (co-) regulation via diverse proposed post-translational modifications in physiological settings. Post-translational regulation of SAMHD1 might not only be achieved by the direct modification of single residues, but also by interaction partners, that could modulate or mediate SAMHD1 anti-viral activity.

A better understanding SAMHD1 regulation in relevant HIV-1 target cells, will also improve our understanding of how SAMHD1 inhibits HIV-1 replication and which conditions license SAMHD1 anti-viral capacity.

## Material and Methods

### Cell lines

Human 293T/17 (ATCC No.: CRL-11268) cells were cultured in DMEM (Sigma-Aldrich) supplemented with 10% fetal calf serum (FCS; Sigma-Aldrich) and 2 mM L-glutamine (Sigma-Aldrich) at 37°C and 5% CO_2_. Human BLaER1 cells (a kind gift of Thomas Graf) (Rapino et al. 2013) cells were grown in RPMI (Sigma-Aldrich) supplemented with 10% FCS and 2 mM L-glutamine at 37°C and 5% CO_2_. For transdifferentiation, 1 x 10^6^ BLaER1 cells per well of a 6-well tissue culture plate were treated with 10 ng/ml human recombinant M-CSF and IL-3 (PeproTech) and 100 nM β-estradiol (Sigma-Aldrich) for 7 days. Half of the cell culture supernatant was replaced with medium containing cytokines and β-estradiol at days 2 and 6. All cell lines were free of mycoplasma contamination, as tested by PCR Mycoplasma Test Kit II (PanReac AppliChem).

### CRISPR/Cas9 knock-out and knock-in

For CRISPR/Cas9 mediated SAMHD1 knock-out (KO), 200 pmol Edit-R Modified Synthetic crRNA targeting *SAMHD1* exon 1 (crSAMHD1_ex1, target sequence: 5’-ATC GCA ACG GGG ACG CTT GG, Dharmacon), 200 pmol Edit-R CRISPR-Cas9 Synthetic tracrRNA (Dharmacon) and 40 pmol Cas9-NLS (QB3 Macrolab) were assembled *in-vitro*, as previously described (Hultquist et al. 2016). Ribonucleoproteins were introduced into 1 x 10^6^ sub-confluent BLaER1 cells using 4D-Nucleofector X Unit and SF Cell line Kit (Lonza), applying program DN-100. Single cell clones were generated using limited dilution one day after nucleofection. To confirm bi-allelic SAMHD1 KO, the modified region was amplified using primer SAM_Seq_Gen-23_FW (5’-GAT TTG AGG ACG ACT GGA CTG C) and SAM_Seq_Gen1116_RV (5’-GTC AAC TGA ACA ACC CCA AGG T) together with GoTaq polymerase (Promega), followed by cloning into pGEM T-easy vector system (Promega) and Sanger sequencing. For knock-in (KI), 100 pmol of the respective ssDNA homologous recombination template with 30 bp homology arms (Dharmacon) to introduce T592A (5’-TAG GAT GGC GAT GTT ATA GCC CCA CTC ATA GCA CCT CAA AAA AAG GAA TGG AAC GAC AGT A) or T592E (5’-TAG GAT GGC GAT GTT ATA GCC CCA CTC ATA GAA CCT CAA AAA AAG GAA TGG AAC GAC AGT AC), as well as Alt-R HDR Donor Oligos (IDT) to introduce H210A (5’-GT GGA ATA AAT CGT CCA TCA AAC ATG TGA GAA AAT GGC CCA GCA CCT TAA AAA CAA AAG CAG CCT TAG AAC AAG AAA AAC ATC), D218A (5’-TC CAT TTC ACC TCC GGG CGA GCA AGT GGA ATA AAT CGT CCA GCG AAC ATG TGA GAA AAT GGC CCA TGA CCT TAA AAA CAA AAG C) or D311A (5’-CA GAA GTG TTC AGT GCA TAC CTG GCA AAA TAA TCC CAT TTG GCG ACG TCA ATG CCA TTT CTT TTA TTA GAT ACT ATC TCA TAA AGG AA) were nucleofected together with ribonucleoprotein complex containing crSAMHD1_ex16 (target sequence: 5’-TTT TTT TGA GGT GTT ATG AG, Dharmacon), crSAMHD1_ex8_1 (target sequence: 5’-TAA AAG AAA TGG CAT TGA TG, Alt-R custom guide IDT), crSAMHD1_ex8_2 (5’-ATG GCA TTG ATG TGG ACA AA), crSAMHD1_ex6_1 (5’-GC TTT TGT TTT TAA GGT CAT GTT TTA GAG CTA TGC T) or crSAMHD1_ex6_2 (5’-CCA TTT TCT CAC ATG TTT GA). To increase KI efficiency, Alt-R HDR Enhancer (V1 or V2, IDT) was added at 1:500 dilution after nucleofection for 24 h. Single cell clones were generated by limited dilution or using Hana single cell dispenser (Namocell) five days after nucleofection. When single cell clones reached confluency, duplicates were generated. One half was lysed (10 min, 65°C; 15 min, 95°C) in lysis buffer (0.2 mg/ml Proteinase K, 1 mM CaCl_2_, 3 mM MgCl_2_, 1 mM EDTA, 1% Triton X-100, 10 mM Tris (pH 7.5)) (Schmid-Burgk et al. 2014) and screened for successful KI using mutation specific custom designed KASP genotyping assays (LGC) and KASP V4.0 2x Master mix (LGC) on a CFX384 Touch Real-Time PCR Detection System (BioRad). Alternatively, mutation specific primer were used to screen for successful KI (H210A 5’-TGA GAA AAT GGC CCA GCA CCT TAA, D218A 5’-TAA ATC GTC CAG CGA ACA TGT GA, D311A 5’-ATA ATC CCA TTT GGC GAC GTC AAT G) using KAPA HiFi HotStart ReadyMix (Roche). Homozygous KI was confirmed by Sanger sequencing after amplification using primer SAM_Seq_Ex16_FW (5’-CAT GAA GGC TCT TCC TGC GTA A) and SAM_Seq_Ex16_RV (5’-ACA AGA GGC GGC TTT ATG TTC C), SAM_Seq_Ex6_FW (5’-GAA TTC AGT TTG GCT GAG TGT GG) and SAM_Seq_Ex6_RV (5’-AAG CAC ATG GGA ATT TTT CAG GAA G), or SAM_Seq_Ex8_FW (5’-TAC AGG CAC TTG CTA CCA TGC CCA AC) and SAM_Seq_Ex8_RV (5’-CTT CTT ATT GCC TCC TCT GGC ACA GC) together with KOD Hot Start DNA Polymerase (Merck) or KAPA HiFi HotStart ReadyMix. Additionally, allele specific sequencing as described for SAMHD1 KO was performed, if required. Absence of large deletions in the region between amplification primers was tested by PCR and analytic gel electrophoresis. Presence of both alleles was confirmed by quantitative genomic PCR (Weisheit et al. 2020), performed using *SAMHD1* exon 16 (FW: 5’-CTG GAT TGA GGA CAG CTA GAA G, RV: 5’-CAG CAT GCG TGT ACA TTC AAA, Probe: /56-FAM/ AAA TCC AAC /Zen/ TCG CCT CCG AGA AGC /3IABkFQ/), exon 6 (FW: 5’-TTT CTT GTT CTA AGG CTG CTT, RV: 5’-AAT ACA TAC CGT CCA TTT CAC C, Probe: /56-FAM/AT TTA TTC C/ZEN/A CTT GCT CGC CCG GA/3IABkFQ/) or exon 8 (FW: 5’-AGG TAC AGC TTC CTT GTT GAA A, RV: 5’-ACA GAC ACG GGC AAA CTT AAT A, Probe: /56-FAM/AG GGA CTG C/Zen/C ATC ATC TTG GAA TCC /3IABkFQ/) specific PrimeTime qPCR Assay (IDT), human TERT TaqMan Copy Number Reference (Thermo Fischer) and PrimeTime Gene Expression Master Mix (IDT) on a CFX384 machine.

### HIV-1 reporter virus infection

VSV-G pseudotyped HIV-1 reporter viruses pNL4.3 E^-^ R^-^ luc (Connor et al. 1995) (HIV-1-luc) and pNL4.3 IRES mCherry E^-^ R^+^ (HIV-1-mCherry) were produced, as detailed previously (Schott et al. 2018). Briefly, pNL4.3 E^-^ R^-^ luc (a kind gift of Nathaniel Landau) or pNL4.3 IRES mCherry E^-^ R^+^ (a kind gift of Frank Kirchhoff) were co-transfected together with pCMV-VSV-G into 293T/17 cells using 18 mM polyethylenimine (Sigma-Aldrich). Filtered (0.45 µm) supernatants were treated with 1 U/ml DNAse I (NEB; 1 h, RT) and purified through a 20% sucrose cushion (2 h, 106750g, 4°C). Viral stocks were titrated for β-galactosidase activity on TZM-bl cells. Virus-like particles containing Vpx (VLP-Vpx) were produced in an analogue manner using pSIV3+ (Nègre et al. 2000) derived from SIVmac251 (a kind gift of Nicolas Manel) and pCMV-VSV-G. Alternatively, pPBj-psi10, VSV-G encoding pMD.G and pcDNA3.1-Vpx (SIVsmm) (Berger et al. 2011; Schüle et al. 2009) were used to produce VLP-Vpx or empty (pcDNA3.1 only) control VLPs. The amount of VLP-Vpx used in all experiments was optimized for complete SAMHD1 degradation. For infection, 3 x 10^4^ cells were seeded per well of a 96-well tissue culture plate. Transdifferentiated BLaER1 cells were allowed to settle for 2 h in medium without cytokines and β-estradiol. VSV-G pseudotyped HIV-1 reporter virus at indicated MOI, as well as VLP-Vpx, were added, followed by spin occulation (1.5 h, 200g, 32°C). For more recent experiments (Fig. 6 and 8) cytokines and β-estradiol were added also after seeding, which increased viability and percentage of CD11b^+^ cell population. Infection was quantified after 24 h by FACS or qPCR (for HIV-1-mCherry), or alternatively by adding 50 µl/well britelite plus reagent (PerkinElmer) and measurement on a Pherastar FS (BMG) (for HIV-1-luc). To show VLP-Vpx mediated SAMHD1 degradation, 4.4 x 10^5^ transdifferentiated BLaER1 cells were treated in a 12-well tissue culture plate in the same conditions and concentrations as stated above.

### Flow Cytometry

For flow cytometric analysis of BLaER1 transdifferentiation and surface marker expression, 1 x 10^6^ native or transdifferentiated BLaER1 cells were collected, washed once in FACS buffer (10% FCS, 0.1% Sodium acetate in PBS; 10 min, 300g, 4°C) and stained with CD11b-APC (M1/70, Biolegend), CD19-PE (HIB19, Biolegend), CD14-PacBlue (M5E2, Biolegend), CD163-PE (GHI/61, BD), CD206-APC (19.2, BD), CD11c-VioBlue (MJ4-27G12, Miltenyi), CD4-APC (RPA-T4, Biolegend), CXCR4-PE (12G5, BD), CCR5-PE (T21/8, Biolegend) or respective isotype controls (Biolegend, BD, Miltenyi) and Fixable Viability Dye eFluor 780 (Thermo Fischer) in presence of FC Block (BD, 20 min, 4°C). Stained cells were washed in FACS buffer twice and fixed in 2% paraformaldehyde (30 min, RT), before analyzing on a LSR II instrument (BD). For readout of HIV-1-mCherry infection, up to six wells of a 96-well plate were pooled and stained with CD11b-APC and Fixable Viability Dye eFluor 780 as detailed above. For intracellular SAMHD1 staining, cells were fixed in Cytofix Buffer (BD; 37°C, 10 min), subsequent to cell surface staining, and permeabilisation with Perm Buffer III (BD; 2min on ice), before staining with anti-SAMHD1 (12586-1-AP, Proteintech) or an isotype control (CST; 60 min, RT) and anti-rabbit IgG-DyLight 405 (Thermo Fischer; 60 min RT). Infected cells were analyzed on a BD LSRFortessa.

### Immunoblot

For immunoblot, cells were washed in PBS, lysed in radioimmunoprecipitation buffer (RIPA; 2 mM EDTA, 1% glycerol, 137 mM NaCl, 1% NP40, 0.1% SDS, 0.5% sodium deoxycholate, 25 mM Tris (pH 8.0)) supplemented with proteinase and phosphatase inhibitor (Roche) for 30 min on ice. Lysate was cleared (30 min, 15000g, 4°C) and protein content was measured by Bradford assay using Protein Assay Dye Reagent Concentrate (BioRad). 20 µg total protein were denaturated (10 min, 70°C) in NuPAGE LDS Sample Buffer and Reducing Reagent (Thermo Fischer) and separated on a NuPAGE 4-12% Bis-Tris gradient gel (Thermo Fischer) in MOPS running buffer (1 M MOPS, 1 M Tris, 69.3 mM SDS, 20.5 mM EDTA Titriplex II). Transfer was performed in an XCell II Blot Module in NuPAGE Transfer Buffer (Thermo Fischer) onto a Hybond P 0.45 PVDF membrane (GE Healthcare). After blocking in 5% BSA or milk powder (Carl Roth) in TBST (Tris-buffered saline, 0.1% Tween; 2 h, 4°C), primary antibodies anti-GAPDH (14C10, CST), anti-Cyclin B1 (4138, CST), anti-Cyclin A2 (4656, CST), anti-SAMHD1 (12586-1-AP, Proteintech), anti-SAMHD1 (A303-691A, Bethyl) and anti-SAMHD1-pT592 (D702M, CST) diluted in 5% BSA or milk powder in TBST were applied overnight at 4°C. Subsequent to washing in TBST, anti-rabbit IgG, horseradish peroxidase (HRP)-linked antibody (CST) was applied (2 h, 4°C) and the membrane was washed again before detection on a FUSION FX7 (Vilber Lourmat) using ECL Prime reagent (GE). If required, membranes were stripped of bound antibody in stripping buffer (2% SDS, 62.5 mM Tris-HCl (pH 6.8), 100 mM β-mercaptoethanol; 1 h, 65°C). Band densities were determined with FUSION software (Vilber Lourmat).

### Quantification of HIV-1 DNA copy number by qPCR

HIV-1-mCherry copy number was quantified by qPCR as detailed previously (Schott et al. 2018). In brief, four wells of CD19-depleted infected, heat inactivated virus (5 min, 95°C) or mock treated BLaER1 cells were harvested and pooled at indicated time points. To reduce background, cells were washed 2 h after infection with medium. Cells were washed and incubated in Proteinase K (Roth) and Ribonuclease A (Roth; 5 min, RT) before isolating cellular and viral DNA with DNeasy Blood & Tissue Kit (Qiagen). qPCR was performed using FastStart Universal Probe Master (Roche) with primers and probes specific for late HIV-1 reverse transcription products (FW: 5’-TGT GTG CCC GTC TGT TGT GT, RV: 5’-GAG TCC TGC GTC GAG AGA TC, Probe: FAM-5’-CAG TGG CGC CCG AAC AGG GA-3’-TAMRA) or reference gene *PBGD* (FW: 5’-AAG GGA TTC ACT CAG GCT CTT TC, RV: 5’-GGC ATG TTC AAG CTC CTT GG, Probe: VIC-5’-CCG GCA GAT TGG AGA GAA AAG CCT GT-3’MGBNFQ) on a CFX384.

### Cellular dNTP levels and concentrations

For measurement of cellular dNTP levels, 2 x 10^6^ transdifferentiated BLaER1 cells were washed in PBS and subjected to methanol extraction of dNTPs, followed by quantification of all four dNTPs by single nucleotide incorporation assay, as described previously (Diamond et al. 2004). CD19 depletion was performed using CD19 microbeads and MS columns (Miltenyi). Cell volumes were determined by seeding respective cell types on a Poly-D-Lysine (Sigma) coated (10%, 1.5h, RT) Cell Carrier-96 well plate (Perkin Elmer). After centrifugation (5 min, 300g), cells were fixed (4% PFA, 15 min, 37°C), permeabilized (0.1% Triton X-100, 5 min, 37°C) and stained using HCS CellMask Deep Red Stain (Thermo Fischer, 30 min, RT). Z-Stack of stained cells was acquired using confocal imaging platform Operetta (Perkin Elmer) and volume was calculated as a sum of cell areas in all relevant Z-stacks using Harmony software (Perkin Elmer).

### Statistical analysis

Statistical analysis was performed using GraphPad Prism (V8). Mean and standard deviations are shown. Statistical significance was assessed using unpaired two-tailed t-test, as well as non-parametric Kruskal-Wallis test or parametric One-Way ANOVA, corrected against multiple testing using Dunn’s or Dunnet correction, respectively.

## Acknowledgements

The authors thank Michaela Neuenkirch and Saskia Mönch for technical assistance, as well as Thomas Graf (Centre for Genomic Regulation Barcelona, Spain) for the BLaER1 cell line, Stefan Bauer (University of Marburg, Germany) for providing BLaER1 cell transdifferentiation protocols and Nadine Laguette (Institute of Molecular Genetics of Montpellier, France) for critical reading and helpful suggestions for the manuscript. In addition, the authors thank Nathaniel R. Landau (NYU School of Medicine, USA) for pNL4.3 E^-^ R^-^ luc, Frank Kirchhoff (University of Ulm, Germany) for pNL4.3 IRES mCherry E^-^ R^+^ and Nicolas Manel (Institut Curie, Paris) for pSIV3^+^ plasmid.

## Author Contributions

Conceptualization, M.S., R.K. methodology and formal analysis, M.S., R.K., A.O., N.V.F.; investigation, M.S., P.R., M.Z., K.S., A.O., N.V.F.; resources, B.K., R.K.; writing—original draft preparation, M.S., R.K.; writing—review and editing, R.K., M.S., B.K., A.O., K.S., P.R., M.Z., N.V.F.; visualization, M.S.; supervision, R.K.; funding acquisition, R.K., B.K.; All authors have read and agreed to the published version of the manuscript.

---

**Figure S1:** Validation of CRISPR/Cas9 mediated mutagenesis of SAMHD1 catalytic core residues. (A) Representative sections of Sanger sequencing traces obtained from genomic *SAMHD1* exon 6 and 8. Base triplets corresponding to modified amino acids are highlighted. Asterisk indicate coding and silent mutations introduced. At least two independent sequencing runs were performed per clone. (B) Quantitative genomic PCR for *SAMHD1* exon 6 and 8 against reference gene *TERT* was performed and 2^-Δct^ value obtained from SAMHD1 KI clones normalized to WT. As a control, half of the WT (WT 1/2) DNA was inoculated and Δct of SAMHD1 calculated against ct of TERT which was obtained in the WT with normal DNA amount. Bar graphs indicate mean of experiments, dots individual biological replicates. Error bars correspond to standard deviation (n = 3).

